# Towards establishing Extracellular Vesicle-associated RNAs as biomarkers for HER2+ breast cancer

**DOI:** 10.1101/2020.09.27.309252

**Authors:** Colin L. Hisey, Petr Tomek, Yohanes N.S. Nursalim, Lawrence W. Chamley, Euphemia Leung

## Abstract

Extracellular vesicles (EVs) are emerging as key players in breast cancer progression and hold immense promise as cancer biomarkers. However, difficulties in obtaining sufficient quantities of EVs for the identification of potential biomarkers hampers progress in this area. To circumvent this obstacle, we cultured BT-474 breast cancer cells in a two chambered bioreactor with CDM-HD serum replacement to significantly improve the yield of cancer cell-associated EVs and eliminate bovine EV contamination. Cancer-relevant mRNAs *BIRC5* (Survivin) and *YBX1* as well as long-noncoding RNAs *HOTAIR, ZFAS1*, and *AGAP2-AS1* were detected in BT-474 EVs by quantitative RT-PCR. Bioinformatics meta-analyses showed that *BIRC5* and *HOTAIR* RNAs were substantially upregulated compared to non-tumour breast tissue, encouraging further studies to explore their usefulness as biomarkers in patient EV samples. We contend that this effective procedure for obtaining large amounts of cancer-specific EVs will accelerate discovery of EV-associated RNA biomarkers for detection of HER2+ breast cancer.

## Introduction

Interactions between tumour and stromal cells sculpt the tumour microenvironment and contribute to cancer malignancy, metastasis and immune evasion. Extracellular vesicles (EVs) [1] mediate one of the key intercellular interactions by shuttling biomolecules in micro and nanoscale lipid-enclosed packages. EVs have been associated in many studies with resistance of cancer to chemo or radio therapies [2].

EVs contain cargo specific to their parental cell, are very stable, and circulate in blood and other bodily fluids. These properties make EVs prime candidates for cancer detection in liquid biopsies [3], either alone or combined with the detection of circulating tumour DNA (ctDNA) or circulating tumour cells (CTCs) [4]. Upregulation of RNA transcripts including long-noncoding RNA (lncRNA) offers a means for distinguishing EVs originating from tumour and non-tumour cells. LncRNAs are greater than 200 nucleotide-long transcripts constituting two thirds of the transcriptome and they appear to play a critical role in carcinogenesis of many cancers including breast malignancies [5]. LncRNAs represent promising EV-associated biomarkers but difficulties in producing sufficient amounts of pure cancer associated EVs complicate validation of lncRNA presence in EVs.

Here, we present a simple solution for obtaining high quantities of cancer-associated EVs by culturing the HER2-positive breast cancer cell line BT-474 in a CELLine AD 1000 two-chamber bioreactor flask. The CELLine bioreactor system mimics physiological growth conditions by allowing 3D cell growth on a fibre-mimetic surface, resulting in increases in cell number as well as EV production [6]. This strategy allowed us to obtain sufficient EV yields to demonstrate that tumour cells release EVs associated with several potential breast cancer biomarkers.

## Methods

### Bioreactor Culture

To prevent bovine EVs present in FCS from contaminating the cancer-specific EVs, we cultured BT-474 cells (seeded at 4.5 × 10^8^ cells/mL) in 15 mL Advanced DMEM/F-12 medium (Gibco, ThermoFisher Scientific, Waltham, USA) supplemented with 2% CDM-HD serum replacement (FiberCell Systems, New Market, USA) in the lower cell chamber of a CELLine AD 1000 bioreactor flask (Argos, Elgin, USA). The same media (150 mL) was used in the upper media chamber but supplemented with 2% fetal calf serum(**Fig 1A**). The dialysis membrane that separates the cell and media compartments allows FCS-specific nutrients < 10 kDa but not EVs to pass through and nourish the cells. Every 3 to 4 days, the 15 mL of conditioned medium from the cell chamber was harvested for EV isolation and the media from the upper chamber was replaced.

**Figure 1.**
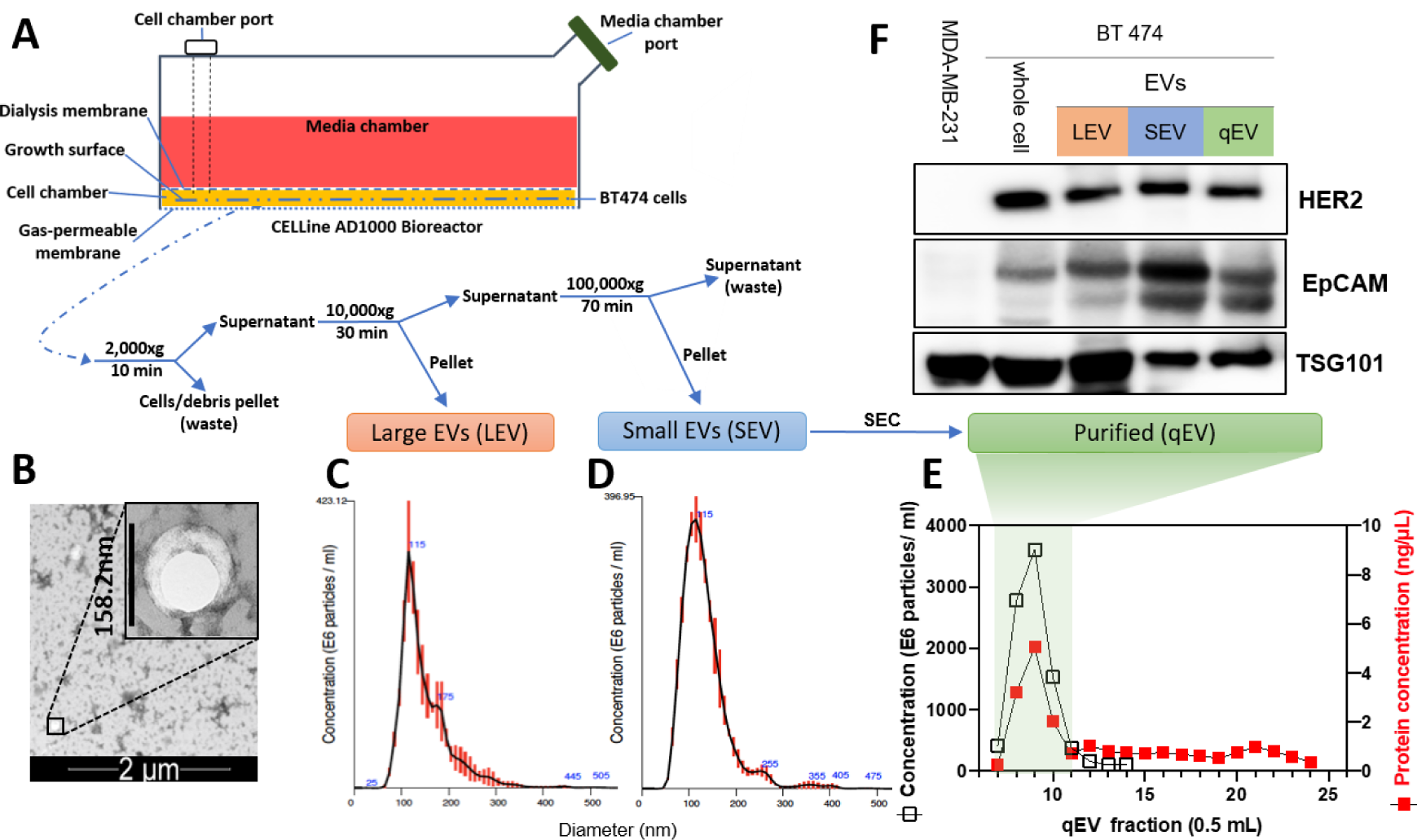
Purification and characterisation of BT-474 EVs. **(A)** experimental procedure employed for EV production, isolation, and purification, **(B)** TEM image of a small EV, **(C**,**D)** hydrodynamic diameter distribution profiles of isolated large and small EVs measured by NTA with red vertical lines and blue numbers denote standard deviation and diameters at specific peaks, respectively, **(E)** EV concentration (empty squares) determined by NTA, and protein levels (filled squares) determined by BCA assay of fractions acquired during separation on a qEV Original SEC column, and **(F)** immunoblot with antibodies specific for HER2, EpCAM and TSG101 proteins. Tetraspanin TSG101 is a loading control. MDA-MB-231 cell lysate serves as the negative control for HER2 and EpCAM proteins. Representative images/data from 3 independent experiments were shown in **B-F**.

### EV Isolation and Purification

EVs were isolated using differential centrifugation and size exclusion chromatography (SEC) as outlined in Figure 1. Conditioned medium (15 mL) was first centrifuged at 2,000 x g for 10 min to remove large debris, 10,000 x g for 30 min to isolate large EVs, and 100,000 x g for 70 min to isolate small EVs (**Fig 1A**). The resulting small EV suspension (in 500 µL PBS) was loaded onto a 35 nm qEV original size exclusion column (Izon, Christchurch, New Zealand), and fractions 7 through 24 were collected using an automated fraction collector (500 µL per fraction). BCA protein quantitation assay (Pierce, ThermoFisher Scientific, Waltham, USA) and Nanosight NS300 nanoparticle tracking analysis (NTA, Malvern Panalytical, Malvern, UK) were performed to quantitate protein and particle concentrations in each fraction, respectively. EV concentrations and size distributions were quantified on NTA by recording three 30 seconds videos under low flow conditions. EV-rich fractions (7-11) were pooled, quantified again using NTA and BCA, and concentrated by ultracentrifugation (Avanti, Beckman Coulter, Brea, USA) at 100,000 x g for 70 min.

### EV visualisation by transmission electron microscopy (TEM)

Negative staining TEM of pooled EV fractions was conducted by adsorption onto Formvar-coated copper grids (Electron Microscopy Sciences, Hatfield, USA) for 2 min, then treating with 2% uranyl acetate for 2 min. Grids were then visualised on a Tecnai G2 Spirit TWIN (FEI, Hillsboro, OR, USA) transmission electron microscope at 120 kV accelerating voltage and images were captured using a Morada digital camera (SIS GmbH, Munster, Germany).

### Protein analysis by Western Blotting

For Western Blot, the proteins (25 μg) were separated by SDS-polyacrylamide gel electrophoresis (PAGE) and transferred to PVDF membranes. Membranes were subsequently immunoblotted with antibodies recognising human HER2 (anti-Neu, Santa Cruz sc-33684), human EpCAM (AbCAM ab223582), and human TSG101 (AbCAM ab30871) and corresponding secondary antibodies. Bound antibodies were visualized using Pierce™ ECL Western Blotting Substrate (ThermoFisher Scientific, Waltham, USA) and the chemiluminescence was measured using a BioRad ChemiDoc MP imaging system (Bio-Rad Laboratories, Inc., Hercules, USA).

### RNA quantitation by qRT-PCR

Technical triplicates of Trizol-purified RNA from each experimental condition were reverse transcribed into cDNA using qScript Flex cDNA kit (Quantabio, Beverly, USA) primed with equal molar ratio of oligo-dT and random primers according to the manufacturer’s instructions. Quantitative RT-PCR was carried out using SYBR Green MasterMix (Life Technologies, Carlsbad, USA) and gene-specific primers previously validated in the literature (Table 1). These included protein-coding mRNAs *EpCAM* [7], *BIRC5* [8], *YBX1* [9], *GAPDH*, and *HPRT1*, and lncRNAs *ZFAS1* [10], *HOTAIR* [11], and *AGAP2-AS1* [12].

**Table 1.**
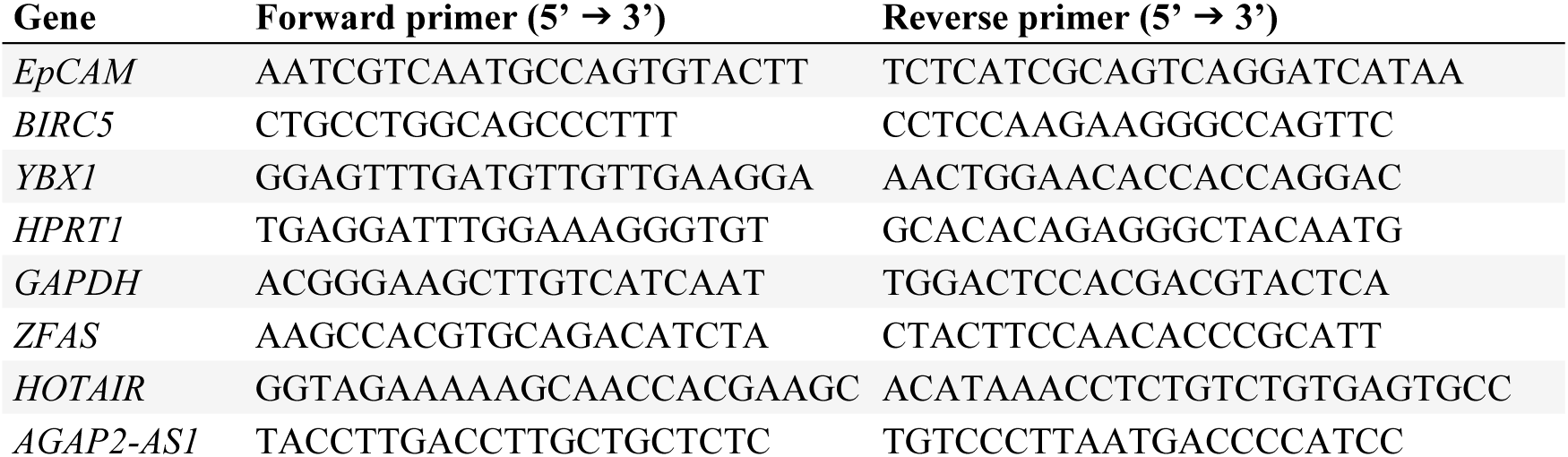
Primers used for quantitative RT-PCR.

### Bioinformatic Meta-analyses

For this meta-analysis, the “RSEM expected count (DESeq2 standardized)” dataset was downloaded on 31st March 2020 from the TCGA_GTEx_TARGET cohort included in the UCSC Xena portal (https://xenabrowser.net/datapages/) and was manually annotated. All data manipulations, plotting and statistical analyses were carried out in R computing environment (v 3.5.3) running in R Studio (v 1.1.414) on a Windows 10 x64 machine. The ggplot2 package (v 3.3.0) was used to render Figures 2B and 2C. Hedges g effect size was calculated using the function cohen.d in the effsize R package (v 0.8.0).

**Figure 2.**
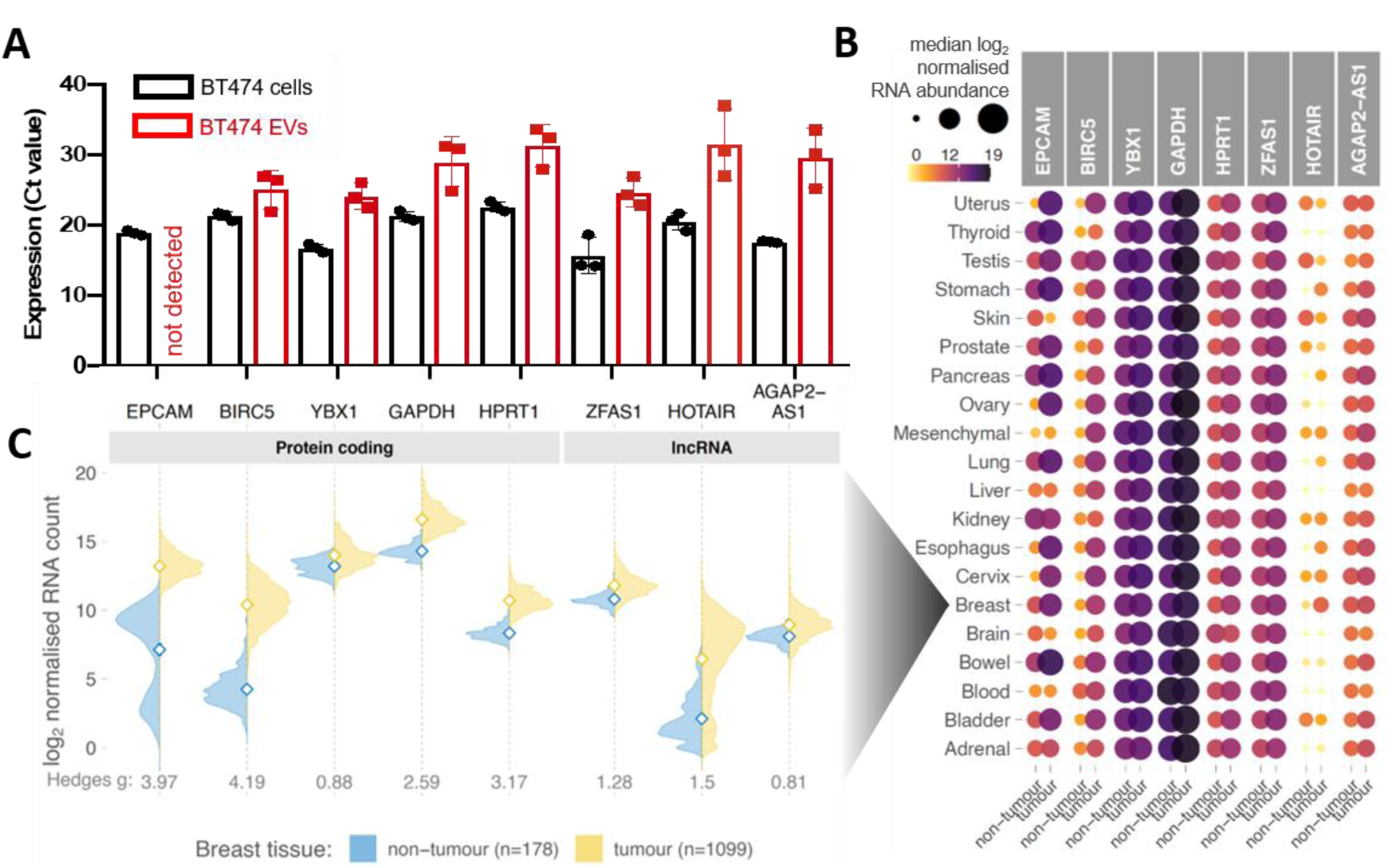
Bioinformatics meta-analysis of BT-474 EV-associated RNAs in tumour and non-tumour tissue. **(A)** Relative mean mRNA abundance of 5 protein-coding genes (*EpCAM, BIRC5, YBX1, GAPDH, HPRT1*) and 3 long non-coding RNAs (*ZFAS1, HOTAIR, AGAP2-AS1*) in BT474 cells and their EVs. Each data point represents the average of three independent experiments (error bars are SEM) **(B)** Comparison of RNA expression of the gene panel studied in (A) between human tumours and their respective non-tumour tissues deposited in TCGA and GTEx portals. Data were manually classified into 20 different organ categories (y-axis) including 8,867 samples across 28 different cancer types and 6,874 samples across 24 non-tumour tissue types. Colour and area of the circles represent median RNA abundance; darker and larger circles indicate higher RNA expression, and **(C)** Distribution of RNA expression of studied genes in breast tumours and breast non-cancer tissues. Open diamonds denote means of each population. Hedges g effect sizes indicate a number of standard deviations that separates the tumour and non-tumour groups. Hedges g > 0.8 demonstrates large effect size, i.e., difference between the means clearly stands out from the “noise” within the groups.

## Results

From three independent experiments, we obtained an average of 1.9 ± 0.3 × 10^11^ large EVs of a mean diameter 150 ± 3 nm and 8.5 ± 0.7 × 10^11^ small EVs of a mean diameter 127 ± 5 nm. Negative-stained transmission electron microscope imaging showed the expected round EV morphology, and NTA size distributions resemble those seen from EVs produced in conventional culture flasks (**Fig 1B-D**). Low levels of contaminating proteins were observed in fractions 11-24 due to 2% CDM-HD serum replacement instead of the standard 5-10% FCS (**Fig 1E**). This allowed the accurate quantification of EV-associated protein markers without the concern of contaminating cellular proteins and demonstrated that the small EVs obtained using ultracentrifugation are suitable for RNA analysis.

Both the BT474 cell lysates and BT474 EVs of all sizes and purities isolated contained TSG101, EpCAM, and HER2 proteins (**Fig 1F**). Consistent with the literature, the triple-negative MDA-MB-231 breast cancer cell line did not express detectable levels of HER2 and EpCAM [13]. TSG101 is a regulator of the endosomal sorting and trafficking process and is expected to be present in both cells and EVs [14]. EpCAM is a cell adhesion glycoprotein which has been used extensively as a liquid biopsy marker for several epithelial cancers [15], whilst HER2 plays an important role in breast cancer subtyping. Interestingly, HER2-positive EVs appear to increase tumour proliferation and resistance to trastuzumab therapy [16].

Quantification of the abundance of several EV-associated RNAs, including protein-coding mRNAs *EpCAM, BIRC5, YBX1, GAPDH, and HPRT*, as well as lncRNAs *ZFAS1, HOTAIR*, and *AGAP2-AS1*, was then performed using qRT-PCR. Despite well-documented differential expression in breast cancer, *EpCAM* mRNA was not found to be associated with the BT-474 EVs, while BT-474 small EVs were clearly associated with established breast cancer-specific RNAs including mRNA *BIRC5* and lncRNA *HOTAIR* (**Fig 2A**).

We then explored the expression of the identical set of RNAs in 15,741 tumour and non-tumour tissue samples included in The Cancer Genome Atlas (TCGA) and Genotype Tissue Expression (GTEx) databases, respectively. Tumour and non-tumour tissues in all 20 tissues analysed expressed similar levels of *YBX1, GAPDH, HPRT1, ZFAS1*, and *AGAP2-AS1* RNAs. This indicates a limited use of these RNAs for differentiating tumour and non-tumour EVs. This result is consistent with the canonical “housekeeping” role of *HPRT1* and *GAPDH* and suggests potential use of *ZFAS1* and *AGAP2-AS1* as housekeeping genes for analyses of lncRNAs in samples including tumour and non-tumour tissues, as well as cultured cells. Of the six candidate biomarkers investigated in this study, only *BIRC5* [8], *EpCAM* [7], and lncRNA *HOTAIR* [11] were found to be differentially expressed in a wide range of cancer types including breast cancer (**Fig 2B-C**).

While EVs hold promise as liquid biopsy targets for breast cancer, efficient production of EVs for molecular characterisation of EV-associated RNA can be challenging using conventional culture systems. In this technical feasibility study, we circumvented this obstacle by culturing BT-474 cells, a commonly used HER2-positive cell line, in a CELLine AD 1000 two-chambered bioreactor which increased the cell density and EV production due to the unique growth surface and fluid interactions [17]. In addition, the common issue of contaminating bovine EVs [18] was avoided by using the serum replacement CDM-HD which is chemically defined, protein free, and animal component free. This bioreactor system provided highly enriched EVs in 15 mL of conditioned media, avoiding the sample loss and extra time associated with pre-centrifugation concentrators. We verified that the EVs contained HER2, EpCAM, and TSG101 proteins. Transmission electron microscope imaging also allowed us to be confident that we had truly isolated small and large EVs in accordance with the MISEV guidelines[19]. We then demonstrated that the BT-474 small EVs were associated with lncRNAs *ZFAS1, HOTAIR*, and *AGAP2-AS*, as well as mRNAs *BIRC5, YBX1, HPRT*, and *GAPDH* using qRT-PCR. Interestingly, the cancer-specific *EpCAM* mRNA was not detected in the small EVs although the EpCAM protein was detectable in the corresponding cell lysates, large EVs, and small EVs. Differential RNA expression in cancer, especially upregulation, has potential to infer a gene’s utility as a biomarker. Our finding indicates that RNAs *BIRC5* and *HOTAIR* are promising EV-biomarkers, particularly in breast cancer, where they are substantially upregulated compared to non-tumour breast tissue.

## Conclusions

Currently, proteins dominate the EV biomarker field. However, novel EV-associated breast cancer biomarkers like lncRNAs and other RNAs are being explored more thoroughly to aid in both detection and management. RNA biomarkers have higher sensitivity and specificity compared to proteins because PCR can amplify traces of RNA sequences with high specificity and sensitivity[20]. Further, it is substantially more economical to detect RNA rather than protein biomarkers because each protein biomarker requires a specific antibody. These findings demonstrate the efficient production of enriched BT474 EVs and highlight their unique cargo, especially *BIRC5* mRNA, encouraging further studies to determine their clinical significance.

## Acknowledgments

The authors would like to acknowledge the support of the Hub for Extracellular Vesicle Investigations (HEVI) and the Auckland Cancer Society Research Centre at the University of Auckland. The authors thank Drs. Bruce Baguley, Graeme Finlay, Marjan Askarian-Amiri, Herah Hansji and Annette Lasham for helpful discussions. EL acknowledges support from the New Zealand Breast Cancer Foundation Belinda Scott Science Fellowship and CH acknowledges support from the New Zealand Breast Cancer Foundation Technology and Innovation Grant. PT acknowledges support of the John Gavin Postdoctoral Fellowship (GOT-1717-JGPDF) from the Cancer Research Trust New Zealand. The funders had no role in the study design, data collection, analysis and interpretation, publication decision or writing of the article.

